# Benchmarking Database Systems for Genomic Selection Implementation

**DOI:** 10.1101/519017

**Authors:** Yaw Nti-Addae, Dave Matthews, Victor Jun Ulat, Raza Syed, Guil-hem Sempéré, Adrien Pétel, Jon Renner, Pierre Larmande, Valentin Guignon, Elizabeth Jones, Kelly Robbins

**Affiliations:** Department of Biotechnology, Cornell University; Boyce Thompson Institute; Centro Internac-ional de Mejoramiento de Maíz y Trigo (CIMMYT); INTERTRYP, Univ Montpellier, CIRAD, IRD; UMR PVBMT, CIRAD; University of Minnesota; UMR DIADE, IRD, University of Montpellier; Bioversity International; College of Agriculture and Life Sciences, Cornell University

## Abstract

**Motivation:** With high-throughput genotyping systems now available, it has become feasible to fully integration genotyping information into breeding programs [22]. To make use of this information effectively requires DNA extraction facilities and marker production facilities that can efficiently deploy the desired set of markers across samples with a rapid turnaround time that allows for selection before crosses needed to be made. In reality, breeders often have a short window of time to make decisions by the time they are able collect all their phenotyping data and receive corresponding genotyping data. This presents a challenge to organize information and utilize them in downstream analyses to support decisions made by breeders. In order to implement genomic selection routinely as part of breeding programs one would need an efficient genotype data storage system. We selected and benchmarked six popular open-source data storage systems, including relational database management and columnar storage systems.

**Results:** We found that data extract times are greatly influenced by the orientation in which genotype data is stored in a system. HDF5 consistently performed best, in part because it can more efficiently work with both orientations of the allele matrix.

**Availability:** http://gobiinx1.bti.cornell.edu:6083/projects/GBM/repos/benchmarking/browse

## 1 Introduction

The development of Next-generation sequencing (NGS) technologies has made it feasible to generate huge volumes of genomic data. In the field of plant breeding, the availability of cost-effective genomic data has the potential to change the way crop breeding is done. Genomic selection (GS) is a breeding method where the performance of new plant varieties is predicted based on genomic information [1]. Multiple studies have shown the potential of this methodology to increase the rates of genetic gain in breeding programs by decreasing generation interval, the time it takes to screen new offspring and identify the best performers for use as parents in the next generation [2], [3]. Although model capabilities exist for the implementation of GS, mainstream applications require the computational infrastructure to manage NGS data, often generated from highly multiplexed sequencing runs that generate low coverage data with large amounts of missing information. The lack of computational infrastructure remains a major barrier to routine use of genomic information in public sector breeding programs.

The Genomic Open-source Breeding Informatics Initiative (GOBii) is a project aimed at increasing the rates of genetic gain in crop breeding programs serving regions in Africa and South Asia by developing the capabilities required for routine use of genomic information. To deal with the technical challenges of storage, rapid access (i.e. query execution), and computation on NGS data, the initial focus of GOBii has been the development of an efficient genomic data management system (GOBii-GDM). The system must be able to efficiently store huge volumes of genomic information and provide rapid data extraction for computation. The system must be scalable for large breeding programs, while being able to run effectively at institutions with limited access to large computational clusters. While many open-source technologies exist for the management of large two-dimensional datasets, it is unclear which technologies best suit the needs of plant breeding and genetics research.

There are many appealing characteristics of traditional relational database management systems (RDBMS), which are designed and built to store, manage, and analyze large-scale data. However, performance can be problematic when dealing with large matrix data like those commonly encountered in genomic research. One common limitation of RDBMS is database partitioning, which allows for a logical database to be divided into constituent parts and distributed over a number of nodes, (e.g. in a computer cluster) [4]. To address these performance issues, many RDBMS have capabilities for working with binary large objects (BLOBs). Current versions of PostgreSQL (version 9.3 and up) have support for JSONB objects that could be used to store BLOBs of genomic data. However, this still does not solve the data retrieval performance issues [4]. An alternative to storing genomic data directly in a RDBMS is to use a hybrid system [5] with high-dimensional data being stored in files and key meta information required for querying the data stored in an RDBMS.

Leveraging several decades of work of the database community on optimizing query processing, columnar store databases, such as MonetDB are designed to provide high performance on complex queries against large databases, such as combining tables with hundreds of columns and millions of rows. In the biological context, experiences show that MonetDB [8] enables the data to be stored in a format to allow fast queries of vectors of genomic data based on marker or sample indexes, which should improve performance relative to RDBMS. More recently NoSQL systems have emerged as effective tools for managing high-dimensional genomic data [9], [10], [11]. NoSQL systems for distributed file storage and searching represent scalable solutions compare to RDBMS when dealing with semi-structured data types [12], [13], and MongoDB, a docu-ment-based NoSQL database has been used to develop a web-based tool for exploring genotypic information [14].

The Hierarchical Data File Format (HDF5) is a member of the high-performance distributed file systems family. It is designed for flexible, efficient I/O and for high-volume and complex data. It has demonstrated superior performance with high-dimensional and highly structured data such as genomic sequencing data [6] making it an appealing option for a hybrid system approach. There are an increasing number of bioinformatics applications, such as BioHDF [20], SnpSeek [7], Oxford Nanopore PoreTools [21] and FAST5, all of which use HDF5 to scale up simple queries execution on large number of documents. However, there is little reported information on the performance of HDF5 when the system is used to process more complex analytical queries that involve aggregations and joins.

To determine the ideal technology to serve as the backend of the GO-Bii-GDM, testing was performed using a large genotype-by-sequencing (GBS) dataset [15], [16], [19]. Open-source RDBMS, PostgreSQL and MariaDB, a community-developed fork under the GNU GPL of MySQL, were used as a baseline for performance testing and compared with HDF5, MonetDB, Elasticsearch [17], Spark [18], and MongoDB. Loading and extraction times were measured using queries that would be commonly run for applications of GS in a breeding program.

## 2 Methods

Six database systems were tested using a subset of a maize nested association mapping (NAM) population GBS SNP dataset [19] containing allele calls for 5258 samples (germplasm lines) and 31,617,212 markers. Each marker represents a physical position in the reference genome where polymorphisms were detected in the samples. Each genotyping call was encoded as a single ASCII character for the diploid result, using IUPAC nucleotide ambiguity codes for heterozygotes and “N” for missing data. The input and output format for all tests was a text file containing only the tab-delimited allele calls.

Genomic matrices can be stored in two different orientations as showing in Figure 1. Given that traditional RDBMS (PostgreSQL and MariaDB) or disk stores are optimized for extracting data by rows, and columnar stores (MonetDB and Parquet) are optimized for extracting data by columns, we stored data in these two different orientations where possible for each of the database systems. Unfortunately, due to size and technology restricts, not all systems could handle both orientations. We tested if the orientation of genotype matrix has an impact on query execution times. For the purposes of this benchmarking, we will define “marker-fast” orientation as the orientation of the genotype matrix in a system that favors the extract of markers, and converse, “sample-fast” as orientation of data in a system that favors the extract of samples. For example, in MonetDB, sample-fast orientation will have samples in columns and markers in rows and as such favor the querying of samples to markers. Vice versa, sample-fast orientation in PostgreSQL will have samples in rows and markers as indexes in a binary JSON object.

**Figure 1:**
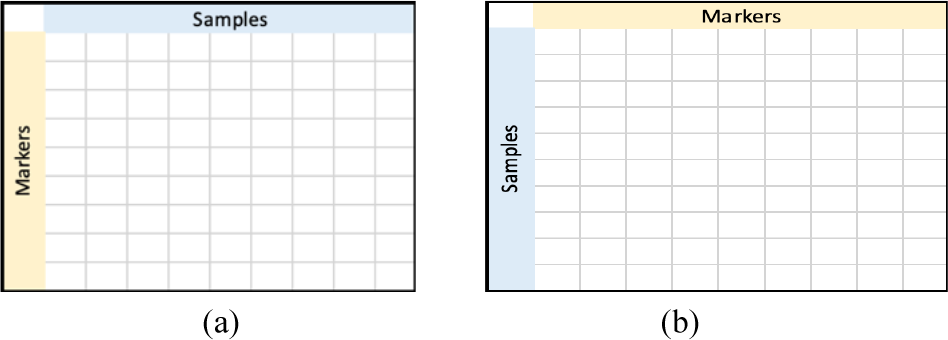
Different orientation of genotyping data. (a) markers in rows and samples in columns, whereas (b) shows markers in columns and samples in rows

Three use cases were used to test the performance of systems with queries set up to extract data by:

I. All samples for a list of markers (USECASE I)
II. All markers for a list of samples (USECASE II)
III. A block of data defined by a list of markers and samples (USECASE III)

For each use case, we tested extracting a contiguous list of markers or samples versus a random list. Care was necessary to avoid unrepeatable timing results due to memory caches in the system. All tests were run on a single node server with a 10 gigabit ethernet connection to a fast-access file server, the server specifications and version of the tested systems are listed in Table 1.

**Table 1.**
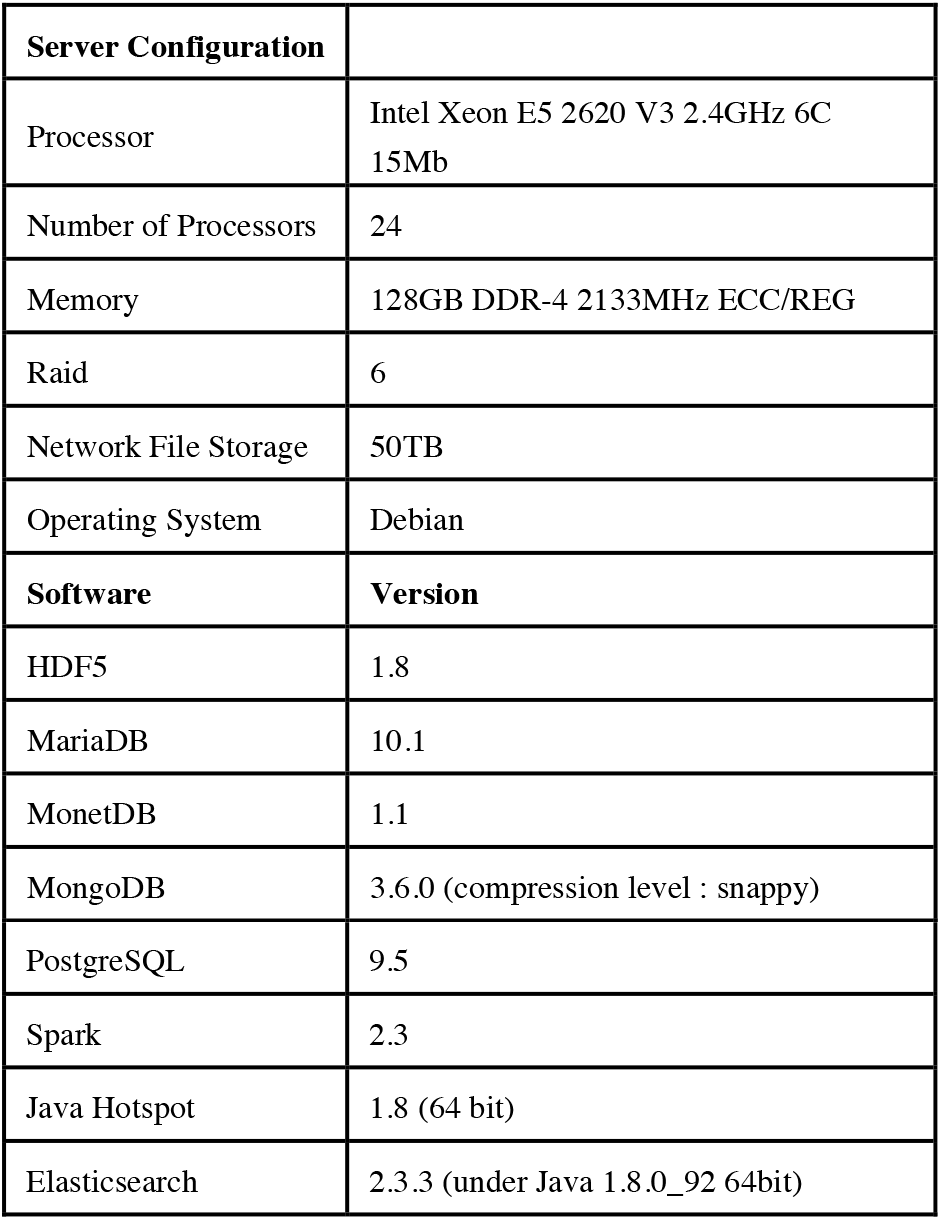
Server configuration and software versions.

| Server Configuration |  |
| --- | --- |
| Processor | Intel Xeon E5 2620 V3 2.4GHz 6C 15Mb |
| Number of Processors | 24 |
| Memory | 128GB DDR-4 2133MHz ECC/REG |
| Raid | 6 |
| Network File Storage | 50TB |
| Operating System | Debian |
| Software | Version |
| HDF5 | 1.8 |
| MariaDB | 10.1 |
| MonetDB | 1.1 |
| MongoDB | 3.6.0 (compression level : snappy) |
| PostgreSQL | 9.5 |
| Spark | 2.3 |
| Java Hotspot | 1.8 (64 bit) |
| Elasticsearch | 2.3.3 (under Java 1.8.0_92 64bit) |

### Database Implementation

The default parameters were applied to each benchmarked system, with some parameters critical for performance, such as memory allocation, manually optimized.

### PostgreSQL Implementation

PostgreSQL is one of the most widely used open source object-relational database systems. Tests where done in version 9.5 and its configuration file was modified to consume up to 30GB of memory per query, whereas the default configurations use only 250MB of memory. Data was stored in “marker-fast” orientation for USECASE I as shown in Table 2.

| Marker | Data* |
| --- | --- |
| S6_120018 | { "B73 (PI550473) : 250028951" : "C", "B97 (PI564682) : 250027958" : "T", "CML103 (Ames27081) : 250027883" : "C", "CML228 (Ames27088) : 250027960" : "C" } |
| S6_120046 | { "B73 (PI550473) : 250028951" : "A", "B97 (PI564682) : 250027958" : "A", "CML103 (Ames27081) : 250027883" : "A", "CML228 (Ames27088) : 250027960" : "A" } |

\*Data stored as JSONB objects

Conversely, data was stored in sample-fast orientation for USECASE II, where samples were in one column and associated markers and SNP allele calls in JSONB format in another column. We recognize that PostgreSQL can be optimized in cases of highly sparse data by storing only non “N” SNPs, which allows PostgreSQL to act like a document store, thereby reducing the size of stored information by many folds. For the purposes of this benchmarking, all data was treated as complete as many USECASEs in genomic selection will pull information in which missing data has been imputed.

### MariaDB Implementation

MariaDB is a community-developed, commercially supported fork of MySQL relational database. MariaDB was configured to utilize up to the maximum available memory on the server. Similar to PostgreSQL, data was stored in marker-fast for USECASE I, as shown in Table 3, and sample-fast for USECASE II. Dynamic columns allow one to store different sets of columns for each row in a table. It works by storing a set of columns in a BLOB and having a small set of functions to manipulate it.

| Marker | Data* |
| --- | --- |
| S6_120018 | { "B73 (PI550473) : 250028951" : "C", "B97 (PI564682) : 250027958" : "T", "CML103 (Ames27081) : 250027883" : "C", "CML228 (Ames27088) : 250027960" : "C" } |
| S6_120046 | { "B73 (PI550473) : 250028951" : "A", "B97 (PI564682) : 250027958" : "A", "CML103 (Ames27081) : 250027883" : "A", "CML228 (Ames27088) : 250027960" : "A" } |

\*Data is stored as dynamic columns

### MongoDB Implementation

MongoDB is an open source document-oriented database system. Mon-goDB tests were performed using version 3.6.0 configured with the WiredTiger storage engine and the snappy compression level, a choice driven by a comparison work done in [14]. MongoDB was tested with data stored in both orientations, marker-fast and sample-fast. For the marker-fast orientation, two types of documents were used:

– one for storing the genotype array corresponding to each marker, as follows:

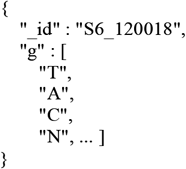

– the second for mapping sample names to indices in the latter array

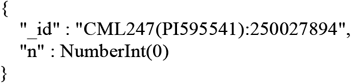

In the sample orientation, the sample collection remained the same as above, but the documents storing genotypes were refactored as:

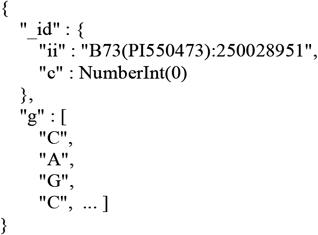

Ideally, the document id would have been just the sample name, and the genotype array length would have been equal to the number of markers, i.e. 31,617,212. But because MongoDB has a 16Mb document-size limitation, we had to split each sample’s genotype array into chunks of maximum size 100,000. This explains why the id here is composite and consists in the following pair: sample name + chunk index.

### HDF5 Implementation

Hierarchical Data Format (HDF5) file format is designed to store and organize extremely large and complex data collections. The allele matrix was stored in one-byte cells in both orientations, marker-fast with samples as columns (HDF5 dimension 1) and markers as rows (dimension 0) and the other in the opposite orientation. When extracting data for a list of samples and a list of markers, USECASE III, the most straightforward approach would be to use HDF5’s *H5Sselect_elements* function to address each allele call by its (sample, marker) coordinates. However, it was found to be much faster to extract all the data for each marker into a memory array using *H5Sselect_hyperslab,* and then look up the results for the desired samples in that array. The speed increase from this approach was more than 30-fold under all conditions tested.

Under some conditions HDF5 performance can be improved by structuring the HDF5 data file in a “chunked” format, depending on the patterns of data access in actual use. A chunk is a contiguous two-dimensional block of the data array that is stored and retrieved as a unit whenever any element within it is accessed. The results reported above were obtained without chunking. Separate tests were performed to compare the unchunked format with four different (marker, sample) chunk dimensions: (256, 256), (1000, 1000), (1000, 5258), and (1, 5258). Loading the HDF5 file was no faster for any of these configurations, and 7-fold slower for (1000, 1000). Retrieval times were also not improved by chunking, using the *H5Sselect_hyperslab* procedure described above.

### Spark Implementation

Spark is an open source distributed general purpose cluster-computing framework. Spark version 2.3.0 was used, using the PySpark (Python 3.6.5) interface. The Java VM used was Java Hotspot 64 bit 1.8.0_171. It is possible there might be a small improvement in benchmark results using the pure Scala interface for Spark, though in general the overhead for using PySpark is not large. Spark benchmarks were run by reading from preprepared Parquet file-format version of the genotype matrix. Parquet is an open source file format for Hadoop and stores nested data structures in a flat columnar format. Besides Spark, Parquet can be used in any Hadoop ecosystem like Hive, Impala and Pig. Data was stored in sample-fast orientation with samples in columns and markers in rows, and vice versa for marker-fast orientation.

### MonetDB Implementation

MonetDB is an open source column-oriented database management system, and version 1.1 of MonetDB was used for the benchmarking. Similar to Parquet file format in Spark, data was stored in sample-fast orientation with samples as columns and markers in rows. Due to number of column restriction in MonetDB, we were not able to store data in marker-fast orientation as the number of markers exceeded the limit for number of columns. MonetDB out of the box is configured to use any available memory on the server.

## 3 Results and Discussion

### 3.1 Data Loading

The load times for each system are presented in Figure 2. HDF5 was the fastest, with MongoDB also performing reasonably well. The two RDMS performed poorly, with MariaDB being the worst, taking approximately 90 times longer than HDF5. While loading time is a lower priority than extraction times, the tight turnaround times from receiving marker data from the lab and generating genomic predictions for selection makes loading times of more than a day for large datasets undesirable for routine implementation of GS. While the process used for loading the data was not optimized, it is unlikely that loading times for MariaDB could be reduced to an acceptable level of performance. The performance of PostgreSQL could potentially be reduced to less than 1 day, but the large gap in performance compared to HDF5, MonetDB, and MongoDB is undesirable. Based on some preliminary testing (data not reported) MongoDB outperformed Elasticsearch (ES). To achieve optimal performance, ES settings must be tuned according to each dataset, hardware environment and expected response time. Furthermore, ES is not designed to return large amounts of data at once, using a scrolling API instead, which complicates the task of gathering query results. Based on these initial results and the complexity of implementing ES as part of GOBii-GDM solution, the decision was made to not pursue further benchmarking on ES.

**Figure 2:**
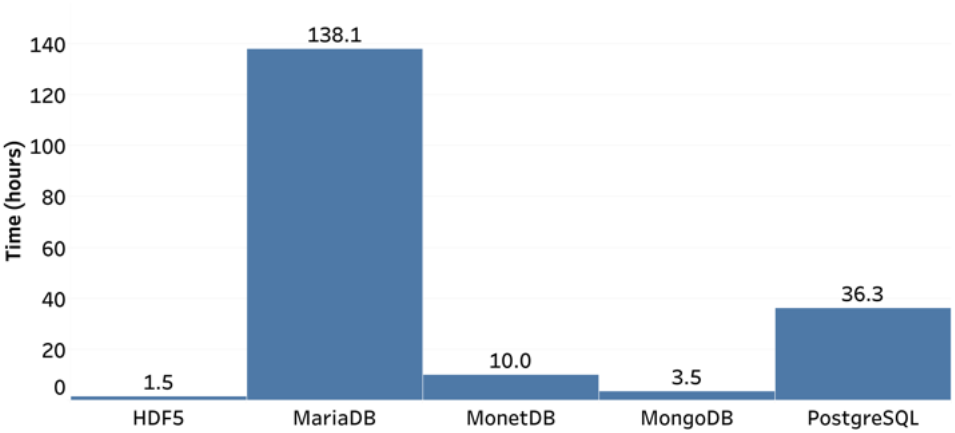
Load times for database systems

### 3.2 Data Extraction

Data extraction tests intentionally extended to very large result sets to test performance beyond the limits expected in actual use. Figure 3 shows the extraction times for increasing number of markers, for all samples in the dataset. For a contiguous block of markers, HDF5 showed the best performance, with the next best solution, MonetDB, performing 11 times slower. On the other hand, for a random list of markers, although HDF5 shows better performance overall, Spark showed a steady performance across different marker blocks, and seems to outperform HDF5 at high marker numbers. The big discrepancy in performance of Spark between contiguous and random list of markers can be explained in the columnar nature of Spark. Spark Parquet file format is a column-oriented data format, so it does not have an “index” of rows and their order, so to ask for a contiguous chunk of rows is antithetical to the design of the data format. Results for MariaDB were greater than 25 hours for all points, off scale in Figure 2a and 2b. When selecting random markers for all samples, MonetDB extraction times exceeded 25 hours for even modest numbers of markers. This is likely due to the orientation of the data stored in the system. Due to limitations in the number of columns for the MonetDB and Spark set-up used for benchmarking, data could not be stored with markers as the columns (> 31 million columns). Given MonetDB is designed for fast querying of columns, the system did not perform well for marker (row) queries.

**Figure 3a:**
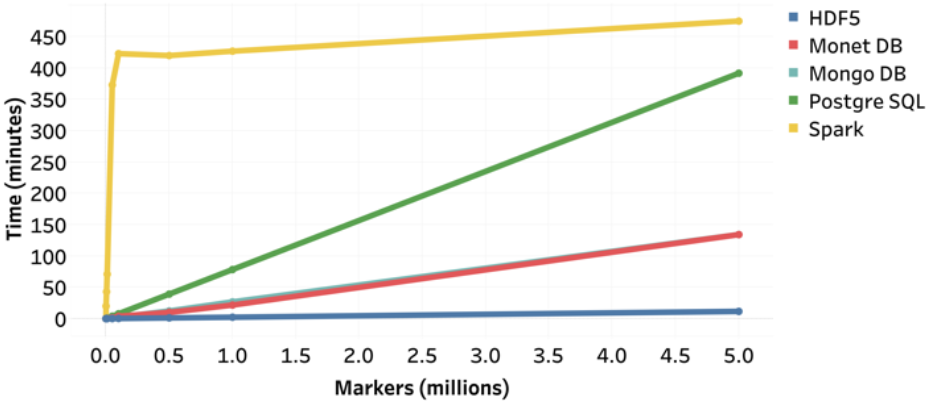
Times for extracting set of contiguous markers for all samples. Times for MongoDB and MonetDB where essentially identical

**Figure 3b:**
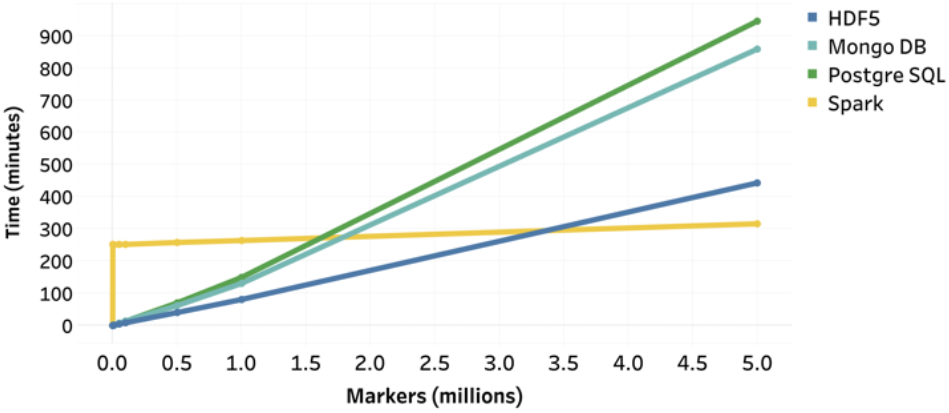
Times for extracting set of random markers for all samples

Extraction times for retrieving all markers for subsets of samples are shown in Figure 3. Again, the results for MariaDB were greater than 25 hours for all queries. As expected MonetDB performed significantly better with queries on samples (columns). As with queries on markers, HDF5 performed best, with the next best solution, Spark, 1.2 and 2.2 times slower than HDF5 for the contiguous and random sample scenarios respectively. For queries on samples, the relative performance of PostgreSQL dropped substantially, with extract times exceeding 20 hours for all data points. The drop in relative performance, and insensitivity to number of samples in the random sample scenario, may be related to the way in which the data is stored in PostgreSQL. A JSONB object is stored for each marker, with each object containing the allele calls for all samples. The observed performance indicates that the total extraction time is influenced more strongly by the time to fetch the JSONB objects from disk into memory, than to extract data for desired samples from the JSONB objects in memory. For the queries extracting all markers from selected samples, two data orientations were tested for MongoDB, with the sample-fast orientation (shown) performing significantly better.

**Figure 4a:**
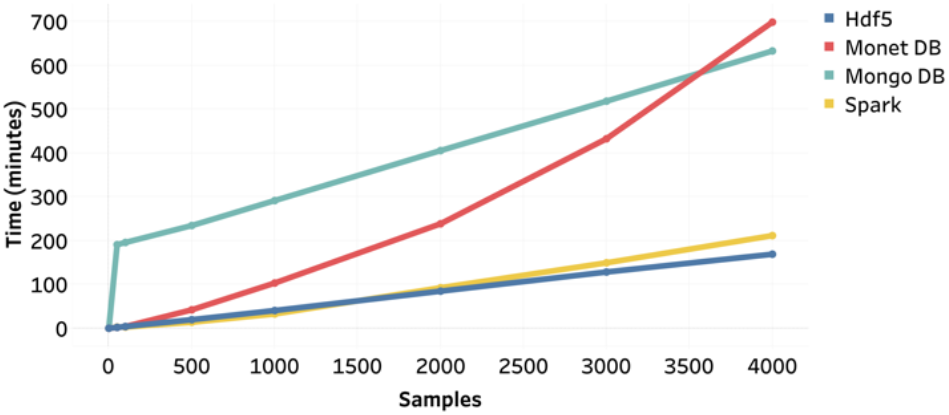
Times for extracting set of contiguous samples for all 32 million markers

**Figure 4b:**
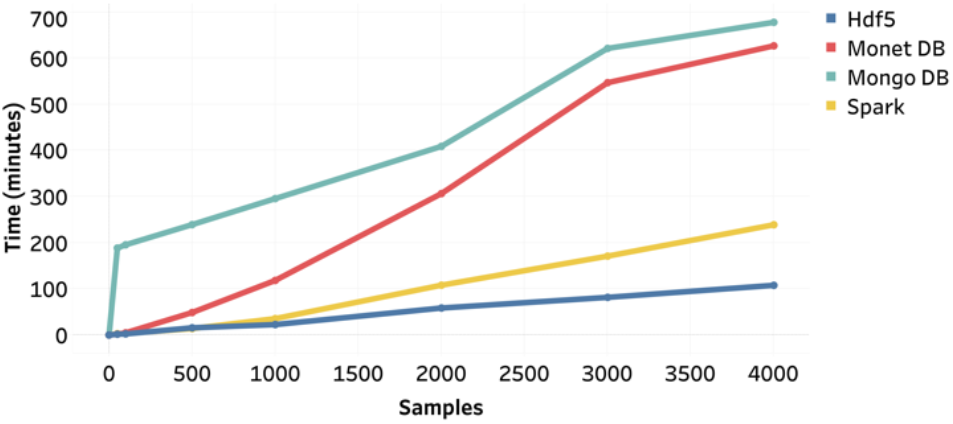
Times for extracting set of random samples for all 32 million markers

Results from USECASE III, extraction based on varying lists of markers and samples. Results for a list of 1 million markers and varying numbers of samples is shown in Figure 5. Once again HDF5 gave the best performance with the next best system performing 5.7 and 1.8 times slower for the largest contiguous and random marker and sample lists, respectively. For the contiguous scenario, MonetDB gave the second-best performance, twice as fast as MongoDB, but exceeded 48 hours for the random list selecting data from 4000 samples and 1 million markers. Neither of the RDBMS performed well for scenario 3, with all extractions taking more than 48 hours for 4000 samples and 1 million markers.

**Figure 5a:**
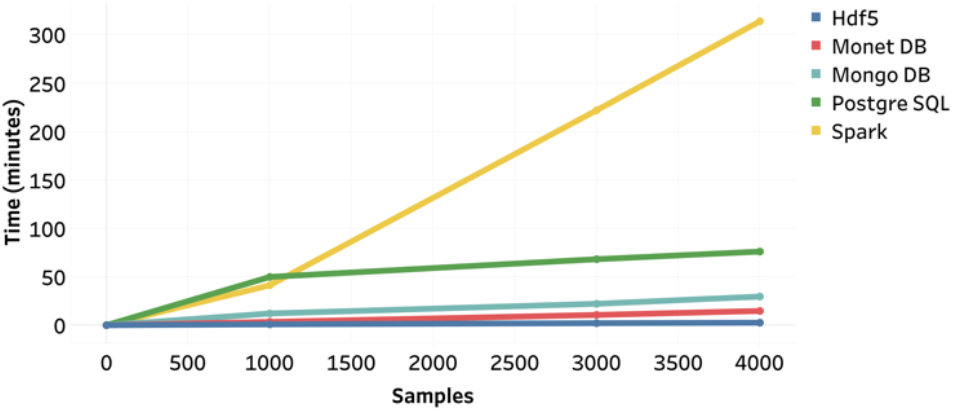
Times for extracting a cross-section of a number of contiguous samples and 1 million contiguous markers

**Figure 5b:**
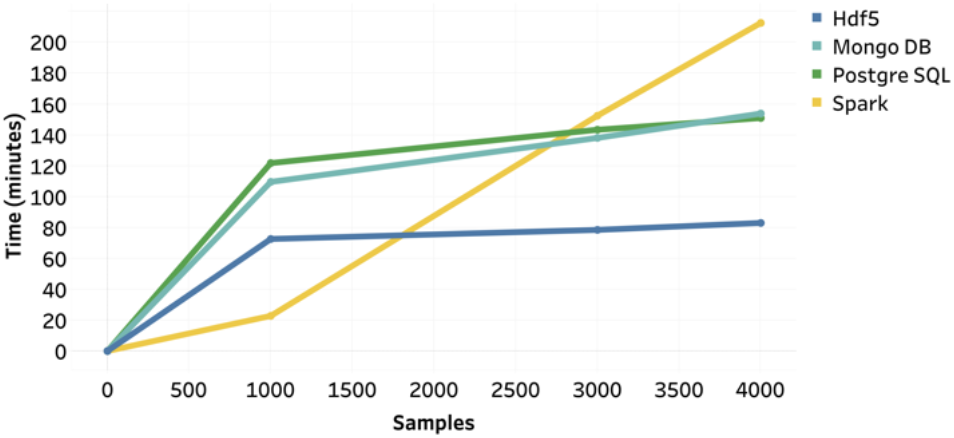
Times for extracting a cross-section of a number of random samples and 1 million random markers

## 4 Conclusion

For all USECASEs the orientation of the data storage had a substantial impact on the performance time of extraction. The fact that PostgreSQL and MonetDB had limitations on storing the data in sample fast and marker fast orientations, reduces their utility for systems that would be regularly queried based on either markers or samples. For systems using HDF5 or MongoDB, best performance would be obtained by storing the data in both orientations with queries being directed to the optimal orientation. While HDF5 showed consistently superior performance in extraction times and loading, implementation in a genomic data management system would require a hybrid approach, with critical meta information likely stored in a RDBMS. A final determination on whether to build a system around HDF5 would need to account for the performance and complexity of developing and deploying a hybrid system. All performance tests were done using a fixed number of cores, but previous studies have shown that the performance of NoSQL distributed file systems, such as MongoDB and Spark, increase with access to more cores [8]. Further testing is required to determine if the performance of MongoDB or Spark would surpass HDF5 when deployed on large clusters.

## Appendix 1

**Table 1:** Results for extracting M markers for all samples

| Con-<br>tigu-<br>ous | # of<br>markers<br>(mil) | MonetD<br>B | HDF5 | Mon-<br>goDB | Post-<br>greSQ<br>L | Mari-<br>aDB | Spark |
| --- | --- | --- | --- | --- | --- | --- | --- |
| Yes | 0 | 0 | 0 | 0 | 0 | 0 | 0 |
| Yes | 0.001 | 0.1 | 0 | 0 | 0.1 | 1566 | 19.5 |
| Yes | 0.005 | 0.2 | 0 | 0.1 | 0.5 | 1623 | 43.45 |
| Yes | 0.01 | 0.3 | 0 | 0.3 | 0.8 | 1523 | 71.3 |
| Yes | 0.05 | 1.2 | 0.1 | 1.4 | 3.9 | 1579 | 373.08 |
| Yes | 0.1 | 3.7 | 0.3 | 2.5 | 7.8 | 1663 | 423.07 |
| Yes | 0.5 | 10 | 1.2 | 12.3 | 39.2 | 1520 | 420.32 |
| Yes | 1 | 21.7 | 2.3 | 26.8 | 78.4 | 1632 | 426.8 |
| Yes | 5 | 134.2 | 11.6 | 134.3 | 391.8 |  | 475 |
| No | 0 | 0 | 0 | 0 | 0 | 0 | 0 |
| No | 0.001 | 44.5 | 0.1 | 0.2 | 0.5 | 1545 | 252 |
| No | 0.005 | 47.2 | 0.4 | 0.7 | 0.6 | 2043 | 251 |
| No | 0.01 | 47.8 | 0.9 | 1.8 | 1.2 | 2108 | 251 |
| No | 0.05 | 58.6 | 4.4 | 7.1 | 6 | 3017 | 252 |
| No | 0.1 | 74.8 | 8.6 | 13.7 | 13.2 | 6021 | 252 |
| No | 1 | 2700 | 80.8 | 130.8 | 149.5 |  | 264 |
| No | 5 |  | 443 | 858.8 | 945.6 |  | 316 |

**Table 2:**
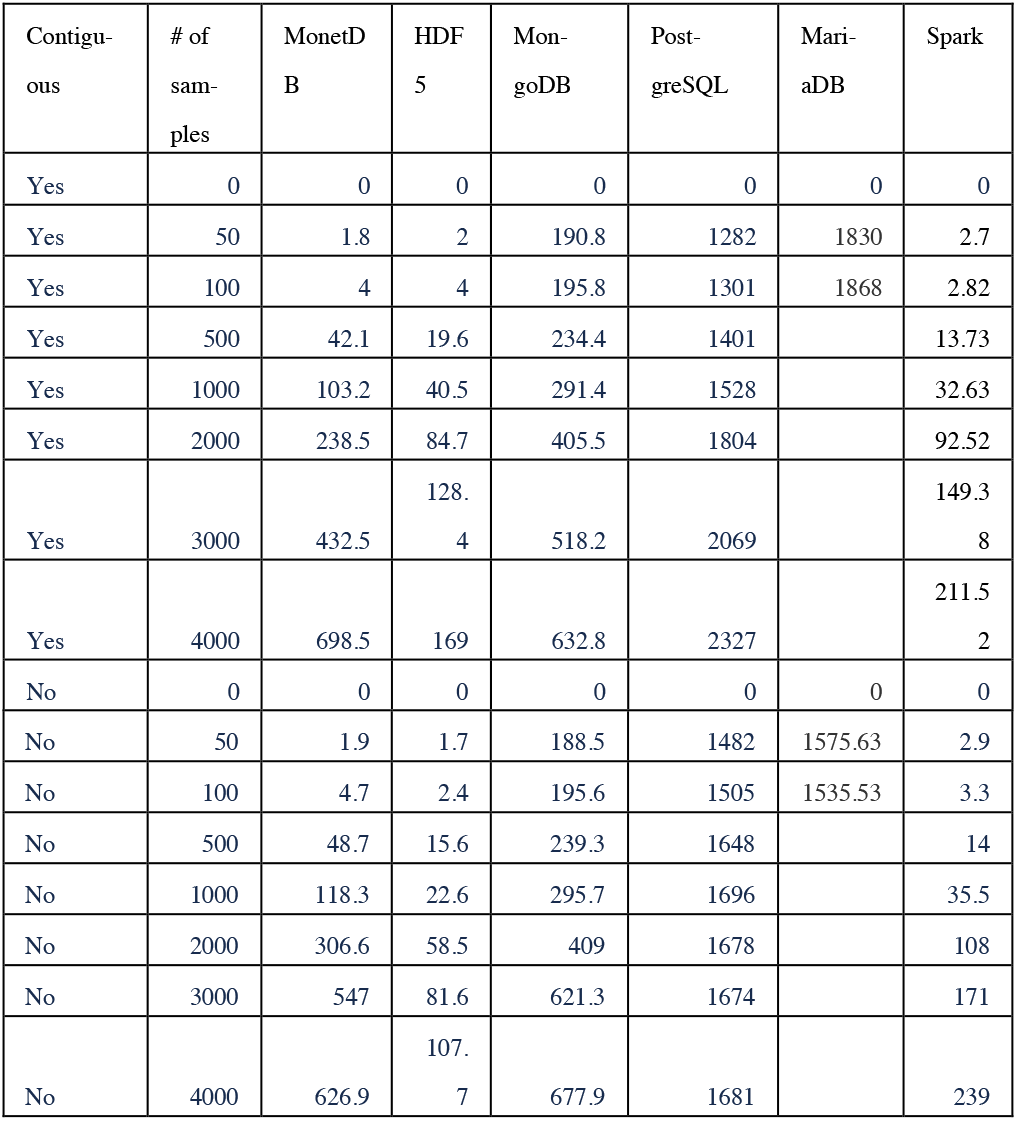
Results for extracting N samples for all markers

**Table 3:**
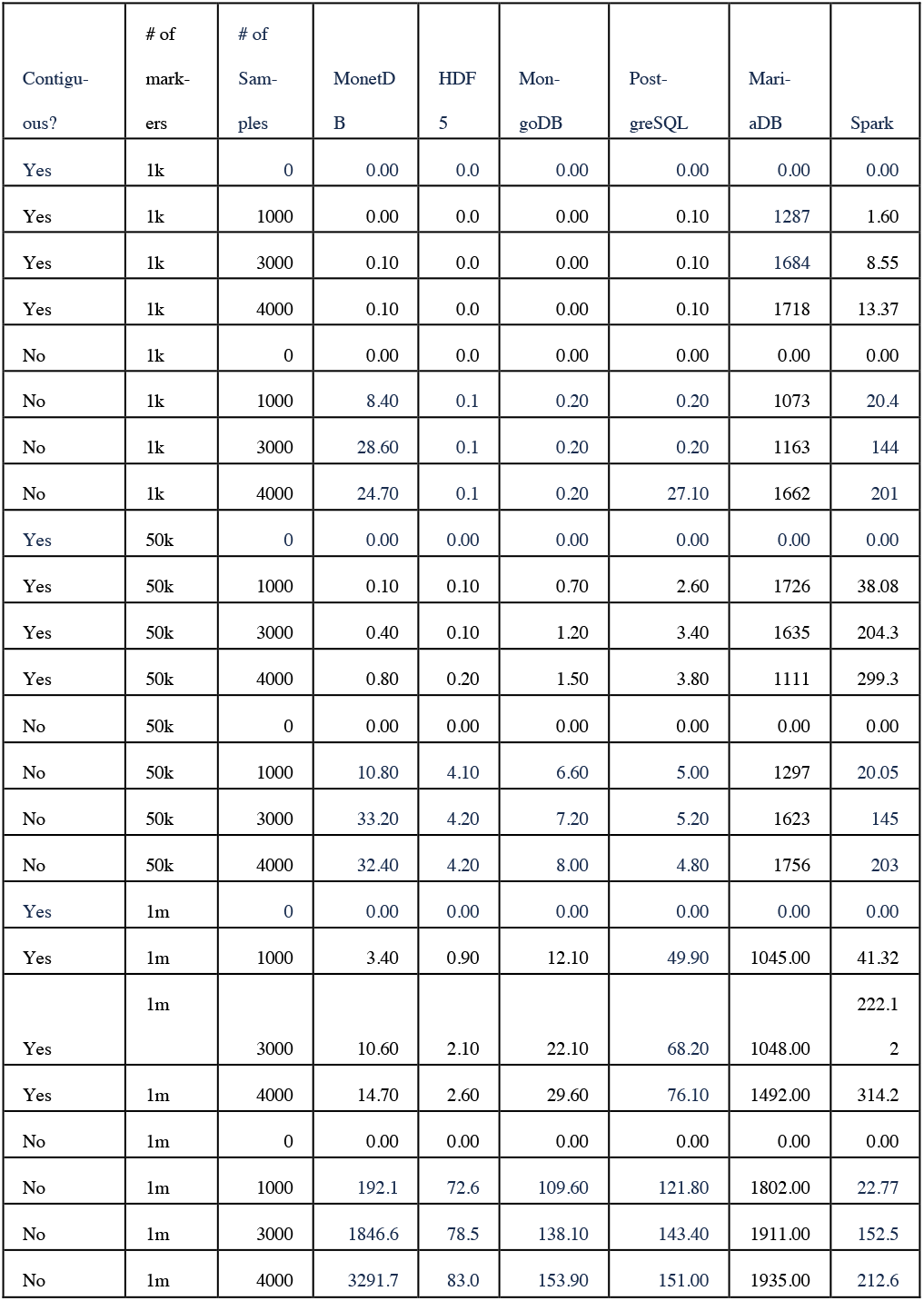
Results for extracting M markers by N samples

